# Excretion initiates walking in the cricket *Gryllus bimaculatus*

**DOI:** 10.1101/362723

**Authors:** Keisuke Naniwa, Yasuhiro Sugimoto, Koichi Osuka, Hitoshi Aonuma

**Author notes:** Correspondence to Hitoshi Aonuma Address: Research Institute for Electronic Science, Hokkaido University, Kita 12, Nishi 7, Sapporo, Hokkaido 060-0812, Japan.

## Abstract

Feces contain information about the donor and potentially attracts both conspecifics and predators and parasites. The excretory system must be coordinated with other behaviors in insects. We found that crickets start walking forward following excretion of feces. Most intact crickets walked around the experimental arena, stopped at a particular site and raised up their body with a slight backward drift to excrete feces. After the feces dropped on the floor, the animal started walking with a random gait pattern away from the feces, and then changed the gait pattern to a tripod gait. Headless cricket also showed walking following excretion. In more than half of excretion events, headless crickets walked backward before excretion. The posture adopted during excretion was similar to that of intact crickets, and post-excretory forward walking was also observed. The occurrence rate of post-excretory walking was more than that of intact crickets. The gait pattern during forward walking was random and never transitioned to a tripod gait in the headless crickets. In animals whose abdominal nerve cords were cut, in any position, pre- or post-excretion walking was not shown in both intact and headless crickets, although they excreted feces. These results indicate that ascending signals from the terminal abdominal ganglion initiate leg movement through the neuronal circuits within thoracic ganglia, and that descending signals from the brain must regulate leg the motor circuit to express the appropriate walking gait.

## Introduction

Internal, as well as external, signals function to initiate stereotyped behaviors in insects. Starvation and thirst can increase motivation to initiate exploratory behavior for food and water, indicating that food digestion and the excretion system must link with initiating particular behaviors in insect. Excretion must be another type of signals to initiate a behavior. The excretory system in insects has been investigated to understand the physiology of food digestion (Maddrell, 1981). Food digestion and excretion in insects are controlled by the peripheral nervous system (Hartenstein, 1997), however, excretion-induced locomotion still remains unclear in insects. Feces contains information about the donor insect. To leave away, or not to leave away after excreting feces, that must be the strategy of insects. Chemicals evaporated from feces potentially attract not only conspecific insects but also predators and parasites. Some insects excrete pheromones with feces. The feces of *Drosophila* adults contains pheromones to attract conspecific flies and increase feeding (Keesey et al., 2016). In locusts, a gregarizing pheromone is also excreted (Obeng-Ofori et al., 1994; Gillett and Phillips, 1977). Increasing population density leads to an accumulates of the excreted gregarizing pheromone, which triggers change in body size, color and behavior of the locust, i.e. solitary phase to gregarious phase (Heifetz et al., 1996). Gregarious locusts have long wings to move long way. Seed harvesting ants mark nesting area with feces, which allows nest mate colony recognition (Grasso et al., 2005). On the other hand, parasitic nematodes are also attracted by the chemicals from insect feces (Schmidt and All, 1979). Therefore, excretion must function as one of the important signals to initiate locomotive behavior in insects.

Here we focus on the excretory behavior of the cricket and found that crickets show walking following excretion with the gait pattern of the after-excretion walking changed from random to tripod. The cricket is solitary insect and live in a burrow to make a territory (Phillips and Konishi, 1973). Leaving from feces position would also prevent from risks of predator and parasites. Cricket change body size smaller, if population density increases (Iba et al., 1995). For crickets, leaving away from feces position would also prevent suffering from density effect. Excretory behavior of the cricket must be a good model system to elucidate the function of feces to initiate adaptive behavior in insects. As a first step in understanding excretion related locomotion in insects, we surgically cut the abdominal nerve cord at different positions. Our results suggest that feedback signals from the hind gut to the abdominal terminal ganglion activates the motor circuits in the thoracic ganglia to initiate walking following excretion in the cricket.

## Materials and Methods

### Animals

Crickets *Gryllus bimaculatus* (DeGeer) were raised in a laboratory colony, reared under a 14h: 10h light and dark cycle (lights on at 6:00 h) at 28 ± 2°C and fed a diet of insect food pellet (Oriental Yeast Co., Tokyo, Japan) and water *ad libitum.* Adult crickets that had molted within 2 weeks before experiments were randomly selected and used in this study.

### Behavior experiments

A cricket was placed in an arena made from clear acrylic sheet (200mm x 300mm x 100mm) to observe the sequences of excretory behavior. The floor of the arena was covered with a matte plastic sheet that was changed after each experiment. Since the inter-excretion interval in a cricket is relatively long, and an intact cricket does not always move around during experiments, we first established a video tracking system to record only the periods when crickets moved (Suppl. 1). We used a web camera (HD Pro Webcam C910, Logicool, Japan) and a Windows 10 based PC to record cricket behavior. The system obtains RGB images continuously at 30 fps and each frame obtained compared with the previous frame. If the image was different from the previous one, the image was saved on PC after a time stamp was inserted into the image. Following recording of long behavior sequences, saved images were converted to MOV format movie files for later analysis. Using this video tracking system, we succeeded in automatically cropping video data where a test cricket was moving.

Recordings were made between 11:00 AM and 9:00 AM of the next day. During recording, the cricket in the arena was illuminated using a LED light source. To investigate excretory behavior, we first drew an ethogram of excretory behavior that focused on direction of movement and gait pattern of walking before and after excretion. We categorized as “resting”, if cricket remained in the same place for more than 10min. We then made a gait diagram to analyze the gait pattern of walking. We measured duration from the time that a leg was placed on the floor. Gait patterns of intact crickets and headless crickets were compared. When a cricket in the arena moved to the corner of the arena where it was difficult to judge the movement of the cricket, we omitted that cricket from the analysis.

### Surgical treatment of the crickets

To elucidate which signals initiate excretory movement, we surgically removed the head of a cricket. Test crickets were anesthetized by exposing them to carbon dioxide gas for 10 s and the head was quickly removed using surgical scissors. The hemolymph from the cut end of the neck was dried, and the headless cricket placed in the experimental arena. After we examined a headless crickets response to air puffs and twitching stimulus of hind leg, we started recording the behavior of the cricket using the video tracking system. In some crickets, the connective of the abdominal nerve cord were cut at different position to elucidate ascending signals from the abdominal ganglion initiate excretory walking.

## Results

### Excretion in the cricket

The excretory behavior of the cricket was stereotyped (Fig. 1) with crickets excreting feces every 3-4 hr (mean ± SD: 197 ± 93 min, N = 7, excretion events (n) = 12). Crickets walked around or remained at the same place in the arena for a while and then stopped moving prior to excretion. When a cricket excreted feces, it raised its body up while slightly moving backwards, and then bent the abdomen towards the ground, so that the excrement dropped on the ground (Fig. 2, Suppl. 2). In about 30% of excretion events in intact crickets, the animals showed backward walking in which they walked a few cycles of steps during which the walking distance was usually short (2.5 ± 1.4 steps, N = 2, n = 5) (Fig. 3).

**Figure 1.**
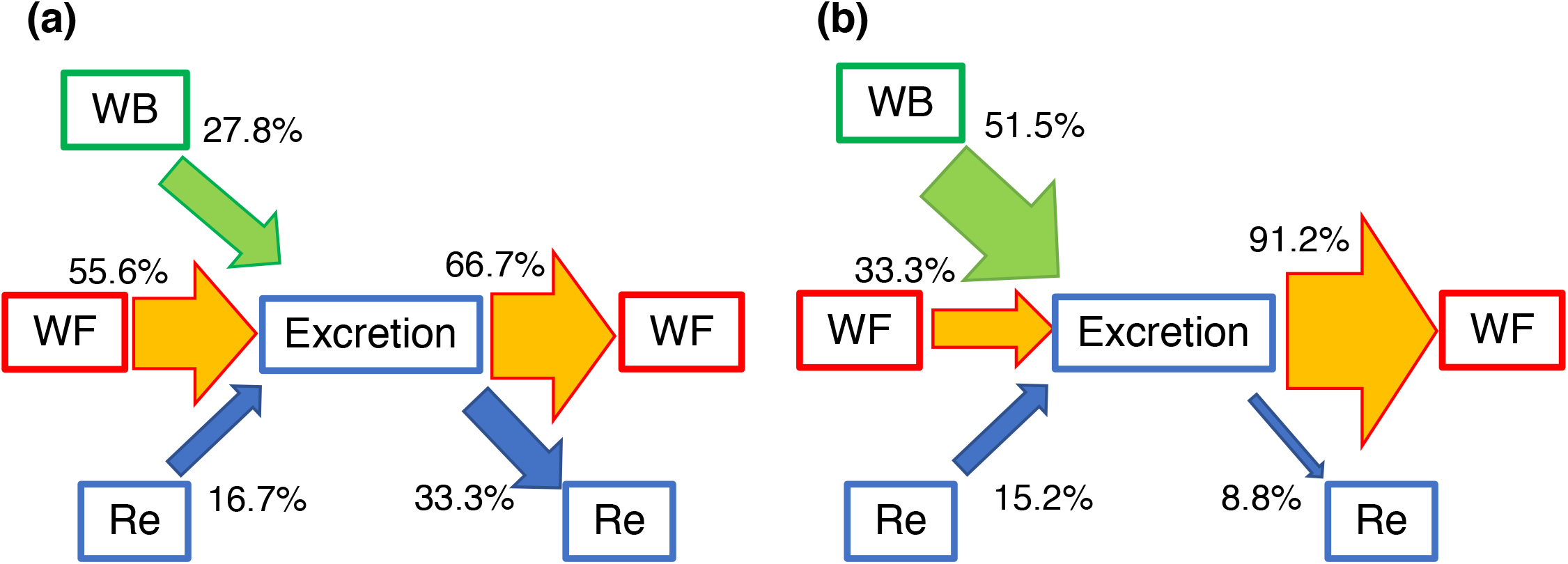
Ethograms for excretory behavior of crickets. These ethograms indicate the behavior of animals before and after excretion. The thickness of each arrow corresponds to the appearance rate. (a) Ethograms for excretory behavior in intact crickets (n=18). (b) Ethograms for excretory behavior in headless crickets (n=33). WB: walking backward, WF: walking forward, Re: resting

**Figure 2.**
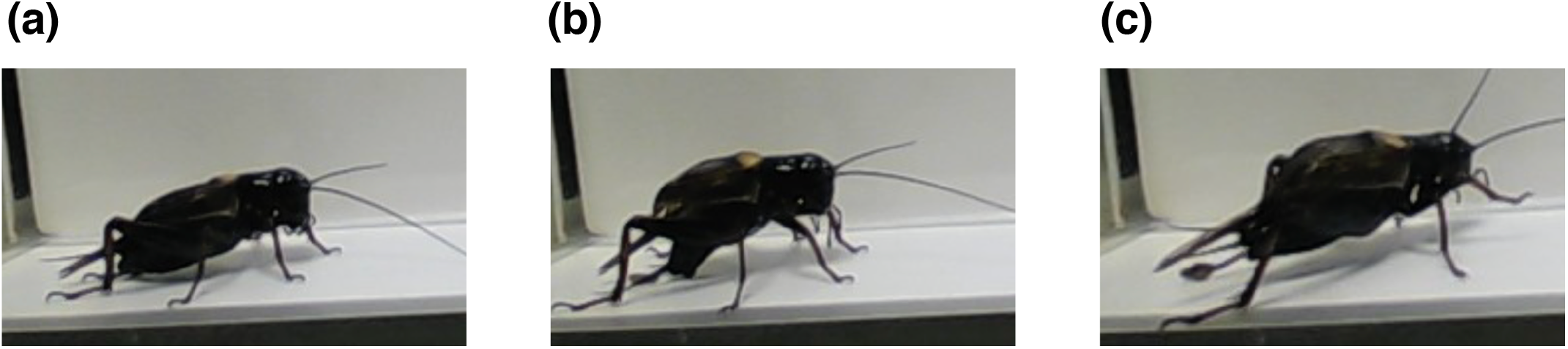
Stereotyped posture of excreting feces of the cricket. (a) A cricket stopped walking. (b) A cricket excreted feces on the ground. (c) After excretion, a cricket start walking.

**Figure 3.**
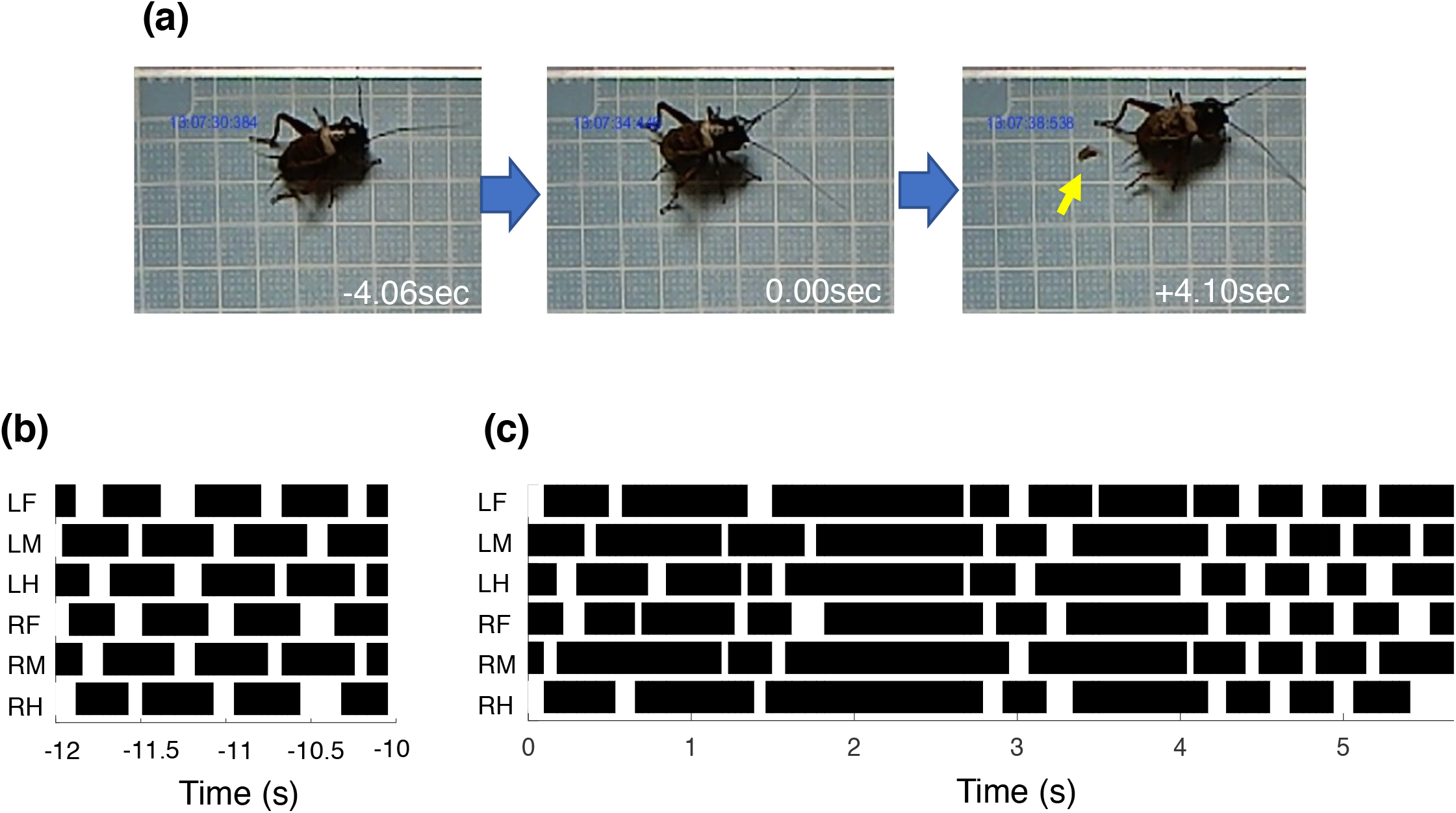
Excretory behavior of an intact cricket. (a) Images of excretory behavior. Crickets stop moving at an appropriate position, raise up and drifted their body backward just before excretion. Some of crickets stepped backward before they raised their body. After excretion, the cricket walked forward to leave the site of excretion. Arrow indicates feces of the cricket. Time 0 s indicates the time of excretion of the feces. (b) Gait diagram of an intact cricket. The cricket walked with a tripod gait before stopping at an excreting site. (c) Gait diagram of an intact cricket just after excretion. The cricket started walking with uncoordinated (random) gait just after excretion. After a few seconds, the gait pattern changed to a tripod gait. L and R in the gait diagrams indicate left and right sides. F, M and H indicate foreleg, middle leg and hind leg respectively.

Excretory behavior was also observed in headless crickets. The inter-excretion interval in headless crickets was similar to that of intact animals (206 ± 147 min, N = 7, excretion events (n) = 26) with no significant differences between intact and headless crickets (Mann-Whitney U test, p = 0.78). The condition of the feces was different between test animals with intact crickets excreting solid feces while headless crickets excreting liquid feces. We found that the backward movement that followed excretion of headless crickets was different from that of intact crickets. Headless crickets showed backward walking following excretion more frequently than intact crickets (p = 0.0008 with U verification) (Fig. 4) in which the number of steps of the hindlegs in backward walking was 13.8 ± 16.5 steps (N = 6, n = 20). The gait pattern of the backward walking in the headless cricket was random (Fig. 4b).

**Figure 4.**
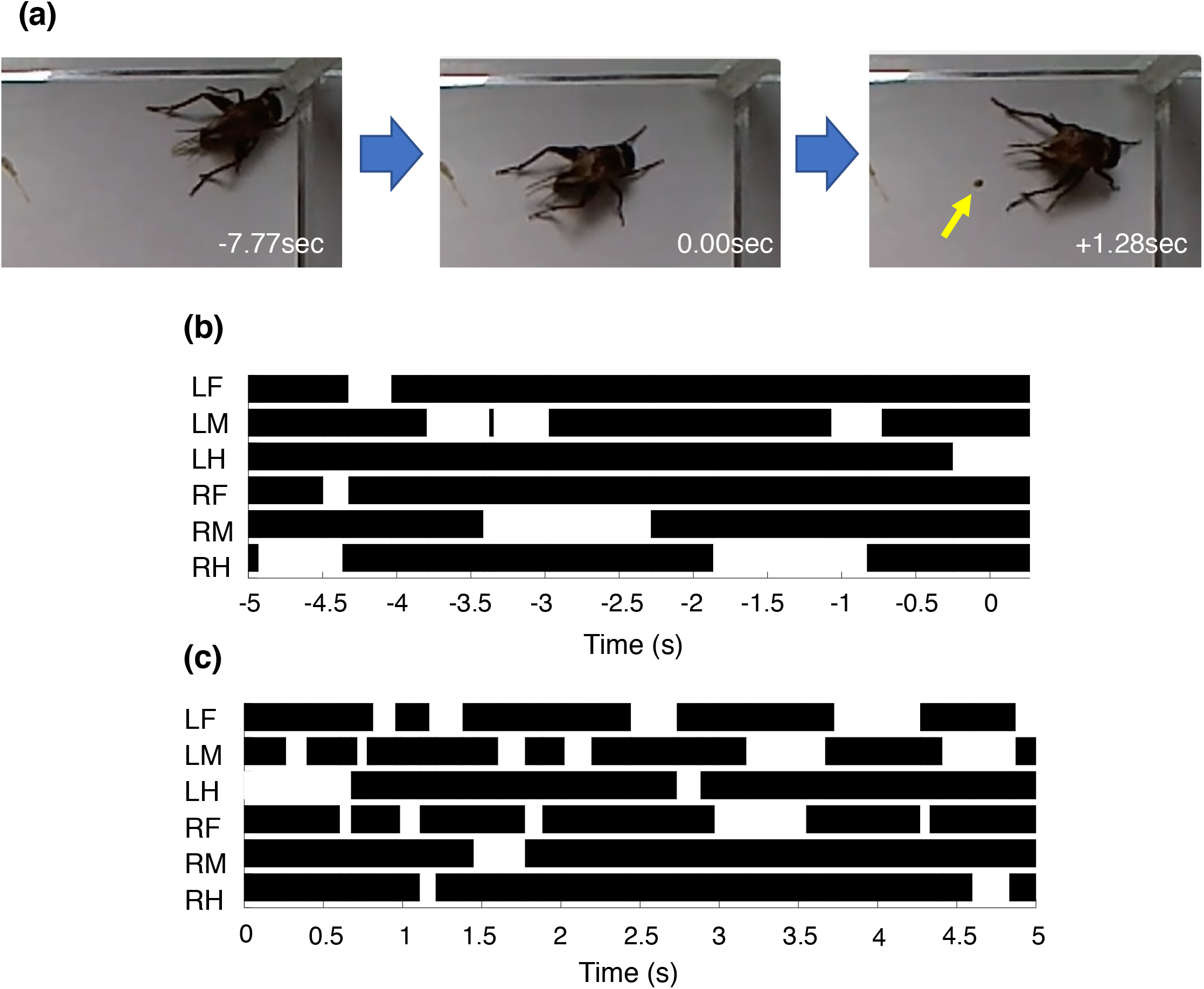
Excretory behavior of a headless cricket. (a) Images of excretory behavior. The headless cricket walked backward and then stopped moving at an excretion site. It raised up and drifted its body backward just before excretion. After excreting feces, it started walking forward away from the excreting site. Arrow indicates feces of the cricket. Time 0 sec the time of excretion of the feces. (b) Gait diagram of the headless cricket. The cricket walked backward with a random gait pattern before stopping at an excreting site. (c) Gait diagram of the headless cricket just after excreting feces. The headless cricket started walking with a random gait until it stopped. L and R in the gait diagrams indicate left and right sides. F, M and H indicate foreleg, middle leg and hind leg respectively.

### After excretion walking in the cricket

Most intact crickets started walking after excreting feces. In about 70%of excretion events (N = 6, n = 12), the cricket walked forward to leave the position of the feces. Approximately 30 *%* of crickets stayed at the same position as excreted (N = 4, n = 6) (Fig. 1a). Intact crickets showed an ordinary tripod gait before excretion (Fig. 3b), however, the gait pattern immediately after excretion was random (Fig. 3c). After a few steps with random gait, the cricket changed its gait pattern to an ordinary tripod gait. By contrast, headless cricket never showed a tripod gait following excretion. In more than 90% of excretion events, headless crickets showed forward walking with random gait until animals stopped walking (Fig. 4c).

To examine if excretion-initiated walking was regulated by the abdominal nervous system, we cut the abdominal nerve cord at different positions and observed the resultant behavior. Crickets whose nerve cord was cut between the 2nd and 3rd abdominal ganglion excreted feces (Table 1), however, they did not show subsequent walking with a random gait pattern. In headless crickets cutting the connectives of the nerve cord between the 2nd and 3rd abdominal ganglion and between the 6th and terminal ganglion resulted in all animals excreting feces. Similar to control crickets, headless crickets did not show backward walking following excretion if the nerve cord was cut at the position between 2nd and 3rd abdominal ganglion. Backward walking just after excretion also disappeared if abdominal nerve cord was cut at a position between the 2nd and 3rd abdominal ganglion and between 6th and terminal abdominal ganglion. For control, we performed sham operation by cutting only cuticle of the abdomen. Headless crickets whose cuticle of abdomen was cut showed backward walking followed by excretion and forward walking after excretion.

**Table 1.**
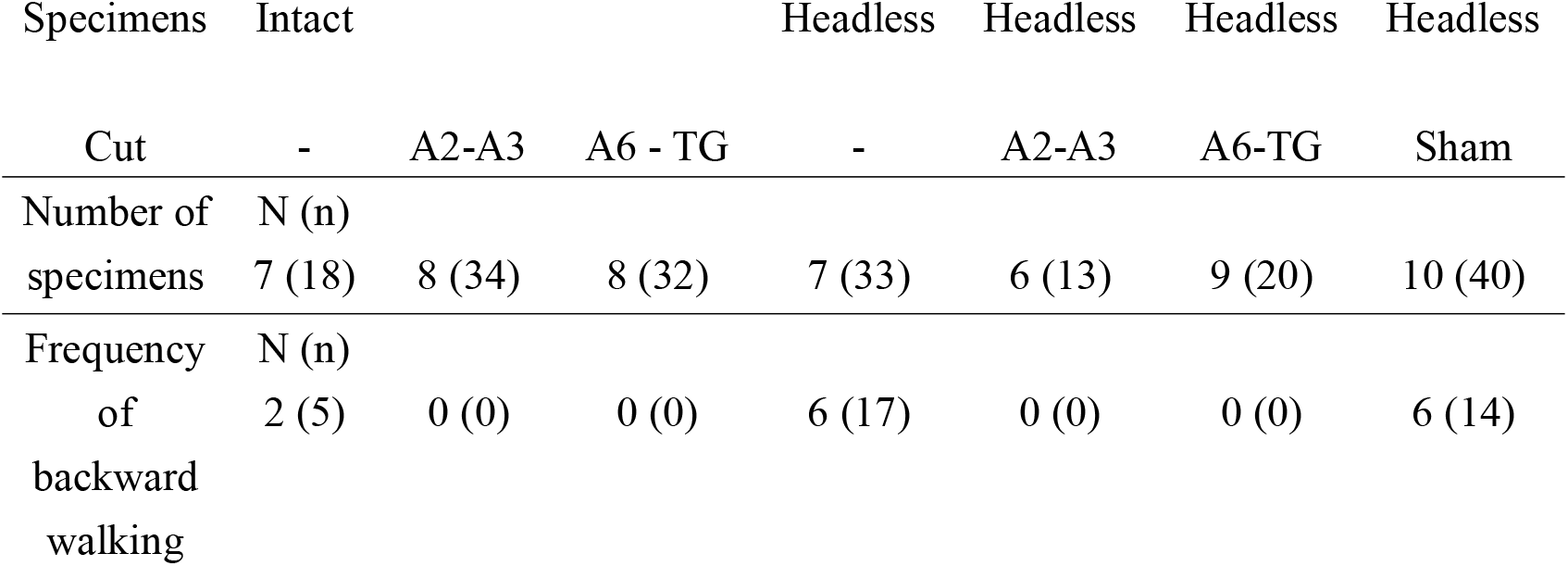
Expression of backward walking followed by excreting of feces in crickets. Backward walking was observed in both intact and headless crickets but not when the abdominal nerve cord was cut between 2nd and 3rd abdominal ganglion, or between the 6th and terminal ganglion. N: number of animals, n: times of excretion, Cut: Position of nerve cord cut, A2-A6: 2nd-6th abdominal ganglion, TG: terminal abdominal ganglion, Sham: sham operation (only cuticle of the abdomen was cut)

## Discussion

This study demonstrates that the excretion initiates leg movements for rapid walking away from feces position in the crickets. The gait pattern immediately after excretion was random and then changed to an ordinary tripod gait. By contrast, headless cricket never showed a tripod gait following excretion.

### Excretion of feces

Cricket showed excretory behavior every 3-4hr in both intact and headless animals in this study. Headless crickets excreted liquid feces suggesting that the rectum absorbs water and that this representative of dysfunctioning excretion system. Removing head ganglia might affect to disturb rectum absorb water in headless crickets. The fecal condition may not affect excretory behavior. Intact and headless crickets whose abdominal nerve cord were cut also excreted feces every 3-4hrs suggesting that the timing of excretion could be controlled by the digestive system in the mid- and hind gut. Indeed, isolated hind guts of the insect are known move rhythmically, and acetylcholine, amino acids and biogenic amines all modulate the movement of the gut (Kanehisa, 1966). Efferent neurons were found innervating the hind gut from the terminal abdominal ganglion in crickets (Hustert and Topel, 1986; Klemm et al., 1986; Elekes et al., 1987; Nässel, 1988). In the cockroach, the sensory afferents that detect movements of the gut innervate the terminal abdominal ganglion (Ishida and Engelmann, 1983). These studies indicate that the information of the movement of hind gut is likely to be processed in the terminal abdominal ganglion.

### Excretion induced movement

We found that headless crickets showed backward walking before excretion and forward walking with random gait pattern after excretion. Although backward walking was not the dominant movement in intact crickets, some of intact crickets showed a short bout of backward walking. When excreting feces, both intact and headless crickets raised their bodies and moved them backwards. The backward walking followed by excreting feces could represent a reflex movement and could be inhibited by descending information from head ganglia including brain and subesophageal ganglion, as in arthropods, information from head ganglia is known to inhibit all reflex activities (Bethe, 1898). Since cricket bent the abdomen to drop feces, the movement of drifting the body backwards could represent a preparatory movement for excretion. To initiate this movement, the cricket need to move the legs so that the body moves backward. Hence, backward walking followed by excretion would be a part of the preparatory movement for an excretory posture.

The backward walking disappeared if the abdominal nerve cord was cut at any position between the 3rd and terminal abdominal ganglion. This indicate that backward walking is initiated by the information from the terminal abdominal ganglion. The afferents from the hind gut must feedback the information of the condition of hind gut to the terminal abdominal ganglion. It was demonstrated that volumetric feedback information detected by hind gut afferents regulates feeding in the locust (Simpson, 1983). Our results suggest that the feedback of information of excretion is processed in the terminal abdominal ganglion and transferred to the thoracic ganglia to initiate leg movements.

Forward walking after excretion was observed in both intact and headless crickets in which the walking could be divided into random gait walks and tripod gait walks. Since headless crickets only expressed random gait walks, tripod gait walking must be coordinated by the descending information from the head ganglia. Most insects show a tripod gait pattern where the foreleg and hind leg on one side with the middle leg of the other side move in synchrony (Wilson, 1966). In the thoracic ganglia, oscillatory neuronal activities of the so-called central pattern generating networks (CPGs) are observed (Borgmann et al., 2009). Descending information from head ganglia is necessary to generate coordinated movement of legs (Heinrich, 2002). Neuronal activity in the central complex of the brain is linked to initiating locomotion in insects (Strausfeld, 1999; Bender et al., 2010). The CPGs are thought to be modulated by descending influences from the brain that initiate, maintain, modify and stop the motor output for walking (Bidaye et al., 2017). The subesophageal ganglion is thought to modulate interactions between sensory inputs from legs and motor output (Knebel et al., 2018). Random gait walks observed after excretion must be a type of reflex activity that is initiated by ascending information from the terminal abdominal ganglion. In intact crickets, ascending information from the terminal abdominal ganglion to the thoracic ganglia initiates walking, and the descending information from the head ganglia coordinates leg movements. Crickets detect air currents using filiform hairs arranged on the surface of cerci of abdomen and respond with quick avoidance movement when they are deflected (Edwards and Palka, 1974). The information of air current is processed and integrated in the terminal abdominal ganglion and the signals transferred to the thoracic ganglia to initiate avoidance walking (Aonuma et al., 2008; Mendenhall and Murphey, 1974; Yono and Aonuma, 2008). This present study demonstrates that feedback information from the hind gut must also acts as a key signal processed in the terminal abdominal ganglion that can initiate walking in crickets.

We conclude that feedback from hind gut that is processed in the terminal abdominal ganglion initiates the sequence of excretory movements in the cricket. The information initiates a backward movement to express an excretory posture and then forward walking. As a next step we now need to investigate the neuronal pathways underlying this sequence of excretory movements, to understand in detail the excretory behavior of insects, and also the neuronal mechanisms underlying walking and gait pattern transition in insects.

## ACKNOWLEDGMENTS

We thank Pro. Philip L. Newland for critical reading our manuscript. This research was supported in part by grants-in-aid for JSPS KAKENHI (Grant-in-Aid for Scientific Research (S), Grant Number JP17H06150) and JST CREST (Grant Number JPMJCR14D5), Japan.

## Author contribution

K.N. and H.A. conceived and designed the experiment; K.N. and Y.S. and H.A. performed the experiment, Y.S. and H.A. analyzed the data; K.N. and Y.S., K.O. and H.A. wrote the paper.

## Supplementary files

**Suppl. 1** Source code of the program for video tracking system.

**Suppl. 2** An intact cricket excreting feces.

**Suppl. 3** Excretory behavior of intact cricket.

**Suppl. 4** Excretory behavior of headless cricket.

